# pH-Dependent Grafting of Cancer Cells with Antigenic Epitopes Promotes Selective Antibody-Mediated Cytotoxicity

**DOI:** 10.1101/785402

**Authors:** Janessa Wehr, Eden L. Sikorski, Elizabeth Bloch, Mary S. Feigman, Noel J. Ferraro, Trevor Baybutt, Adam E. Snook, Marcos M. Pires, Damien Thévenin

## Abstract

A growing class of immunotherapeutic agents work by redirecting components of the immune system to recognize specific markers on the surface of cancer cells and initiate a selective immune response. However, such immunotherapeutic modalities will remain confined to a relatively small subgroup of patients until two major hurdles are overcome: (1) the specific targeting of cancer cells relative to healthy cells, and (2) the lack of common targetable tumor biomarkers among all patients. Here, we designed a unique class of agents that exploit the inherent acidic microenvironment of solid tumors to selectively graft the surface of cancer cells with immuno-engager epitopes for directed destruction by components of the immune system. Specifically, conjugates were assembled using an antigen that recruit antibodies present in human serum, and the pH(Low) Insertion Peptide (pHLIP), a unique peptide that selectively target tumors *in vivo* by anchoring onto cancer cell surfaces in a pH-dependent manner. We established that conjugates can recruit antibodies from human serum to the surface of cancer cells, and induce complement-dependent and antibody-dependent cellular cytotoxicity by peripheral blood mononuclear cells and also an engineered NK cell line. These results suggest that these agents have the potential to be applicable to treating a wide range of solid tumors and to circumvent the problem of narrow windows of selectivity.

## Introduction

Recent advances in targeted immuno-oncological treatments are starting to reveal the tremendous power in deploying the immune system against cancer cells. A subclass of these agents operate by directing antibodies or immune cells to attack specific markers on the surface of cancer cells. It includes clinically approved treatments such as Rituximan (Rituxan®), Obinutuzumab (Gazyva®) and CAR-T cells and also synthetic agents that re-engage the immune system by redirecting antibodies to cancer cell surfaces. The latter are agents often based on bifunctional molecules composed of: a tumor-homing moiety that targets over-expressed membrane proteins (e.g., folic acid receptor, prostate specific membrane antigen) and an immunogenic epitope such as 2,4-dinitrophenol (DNP) (*1*) and α-galactose (*2, 3*) that can recruit antibodies circulating in human serum. Decoration of cancer cells with antigens leads to the recruitment of antibodies (opsonization) and subsequent killing of cancer cells through complement-dependent cytotoxicity (CDC) or antibody-dependent cellular cytotoxicity (ADCC) pathways. These agents have shown exciting anti-cancer activities *in vitro* and tumor reduction *in vivo* (*1, 3-15*).

A significant drawback to prior approaches is the reliance on small changes in expression levels of surface biomarkers for targeted surface grafting of antigens. As a direct consequence of this targeting modality, it is expected that there is wide patient response levels (due to variable biomarker distribution and expression across patient populations), and development of acquired resistance and relapse (due to loss or mutation of biomarker) (*16-18*). Indeed, cancer biomarkers targeted by monoclonal antibodies or small-molecules tend to be over-expressed in a tumor-associated, not tumor-specific, manner (*19, 20*). Moreover, these strategies are not only hampered by variable antigen distribution across patient populations, but also ineffective against cancer types lacking specific biomarkers (e.g., triple-negative breast cancer). Thus, reliance on surface protein biomarkers as the basis for targeted immunotherapy has clear limitations and will invariably result in the rapid selection of drug-resistant tumor clones that lose biomarker expression.

In contrast, we developed a new and unique class of pH-sensitive bifunctional agents capable of selectively decorating the surface of cancer cells with antigens and of promoting directed antibody-mediated cytotoxicity against cancer cells. Our tumor-targeting and tagging approach is based on the pH(Low) Insertion Peptide (pHLIP), a family of peptides with established selective tumor targeting and exciting therapeutic potential that anchor onto the surface of cancer cells in a pH-dependent manner. This strategy takes advantage of a major vulnerability of solid tumors: the inherent acidity of their microenvironment. Indeed, unlike healthy tissues, the microenvironment surrounding nearly all tumor masses regardless of their tissue or cellular origin is acidic (pH 6.0-6.8) (*21-26*), providing a window for selective targeting tumors. Notedly, pHLIP displays unique properties that set it apart from other pH-sensitive agents: (i) its physico-chemical properties can be readily tuned (*27-33*), (ii) because its insertion is unidirectional (i.e., extracellular N-terminus), pHLIP can be used to graft a variety of molecules conjugated to its *N*-terminus onto cancer cell surfaces (*34-36*), (iii) it accumulates in neoplastic tissues while avoiding healthy tissues in many types of human, mouse, and rat tumors including breast cancer and metastatic lesions (*29, 37-40*), and (iv), unlike other pH-sensitive strategies that rely solely on the bulk pH surrounding the tumor surrounding, pHLIP peptides operate at the surface of cancer cells where the pH is the lowest (another 0.3-0.7 pH units lower than bulk) (*41*). This provides pHLIP with superior cell surface residency times and tumor penetration. Here, we show that pHLIP peptides can selectively decorate the surface of cancer cells with a clinically relevant antigen, resulting in a pH-dependent recruitment of antibodies and targeted cancer cell killing by a combination of CDC and ADCC.

## Results

### Design and Synthesis of DNP-pHLIP conjugates

The design of our bifunctional agents consists of the model immunogenic 2,4-dinitrophenyl (DNP) moiety and a panel of pHLIP sequence variants (DNP-pHLIP). DNP is used as a proof-of-principle antigen because of (i) the presence of anti-DNP antibodies in most human serum (*42*), (ii) its synthetic tractability in assembling conjugates, and (iii) its demonstrated potential to provoke an immune-based clearance of the tumors *in vivo* (*1, 43*). Peptides in the pHLIP family are soluble in neutral aqueous solutions as monomers but adopt an inducible transmembrane α-helix under acidic conditions. With the goal of tuning the insertion parameters, three pHLIP variants with established properties were chosen: pHLIP(WT), pHLIP(D25E) and pHLIP(D14Gla,D25Aad) (see Methods). Both pHLIP(D25E) and pHLIP(D14Gla,D25Aad) have pH_50_ (i.e., the pH at which 50% of pHLIP peptides are in the α-helical membrane inserted state) higher than of pHLIP(WT): 6.1 vs. 6.5 and 6.8, respectively (*27, 32*). Indeed, we hypothesized that higher pH_50_ of insertion should result in greater overall pHLIP-based surface tagging. At first, three DNP-pHLIP conjugates were prepared by standard Fmoc solid-phase synthesis incorporating an *N*-terminal lysine residue modified with a DNP substituent (L-Lys(DNP)-OH) (see Methods). DNP-pHLIP conjugates were purified by RP-HPLC and characterized by MALDI-TOF mass spectrometry (Figure S1). Since pHLIP insertion in lipid membrane is unidirectional (i.e., *N*-terminus oriented towards the acidic environment), placement of the antigen at the *N*-terminus should result in cancer cell surfaces decorated with DNP epitopes facing the extracellular environment, thereby engaging with anti-DNP antibodies from serum (Figure 1). Importantly, we showed that conjugation of DNP to pHLIP variants does not significantly affect their respective pH-mediated insertion into the membrane of large unilamellar POPC lipid vesicles, as monitored by circular dichroism spectroscopy (Figure S2). Indeed, all three conjugates exhibit a characteristic transition to an α-helical structure when the pH is decreased.

**Figure 1.**
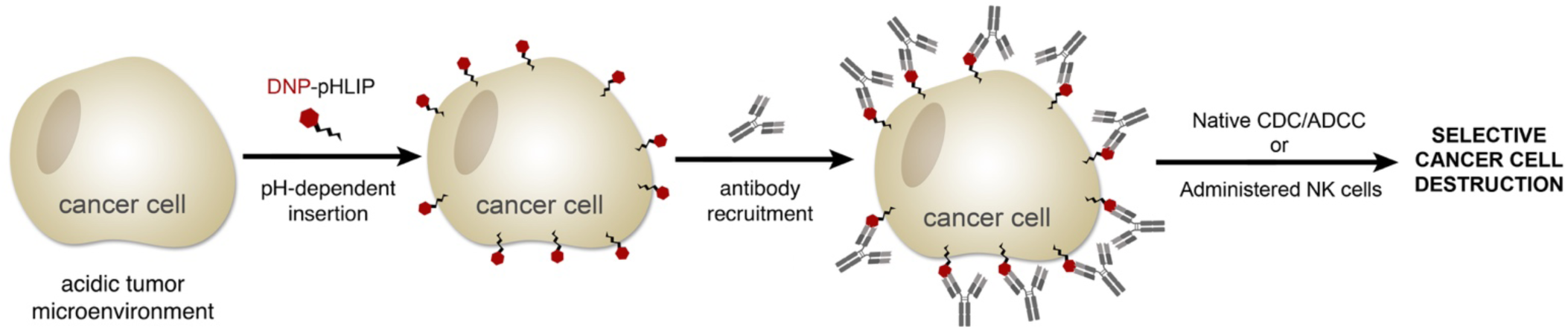
Proposed mechanism of DNP-pHLIP conjugates. Tumor cell surfaces are selectively decorated with the DNP antigen that direct the selective recruitment of endogenous antibodies, in turn engaging and activating both complement and cellular components of the immune system against cancer cells.

### DNP-pHLIP conjugates tigger opsonization of cancer cells in a pH-dependent manner

We first evaluated the potential of DNP-pHLIP conjugates to promote the selective recruitment of anti-DNP antibodies to the surface of triple-negative breast cancer MDA-MB-231 cells. Briefly, cells were treated with DNP-pHLIP conjugates at pH 7.4 or 6.0 (to mimic tumor microenvironments) and incubated with anti-DNP IgG antibody labeled wi fth AlexaFluor-488 (anti-DNP-A488). The relative amount of bound anti-DNP-A488 was quantified by flow cytometry, which should reflect the level of DNP epitopes available for engagement with their cognate antibodies. The concentration of anti-DNP IgG antibodies used here (25 µg/mL) was on par or even below that found in human serum (*42, 44, 45*). Satisfactorily, we observed a pH-dependent increase in cellular fluorescence levels for all three conjugates (Figure 2A), suggesting that DNP is displayed on the cancer cell surface in an orientation that promotes antibody binding. Crucially, no increase in cellular fluorescence was observed when cells were treated with DNP-pHLIP conjugates followed by incubation with a mock FITC-labeled anti-human IgG (instead of anti-DNP-A488 antibodies, Figure S3). These results indicate that the increase in fluorescence signal observed with anti-DNP antibodies is due to the specific recruitment of antibodies by DNP-pHLIP conjugates, and not nonspecific antibody absorption onto cell surfaces. The level of presented DNP epitopes on the cancer cell surface also appears to remain stable over time based on findings that cellular fluorescence levels remained relatively constant over 1-hour incubation (Figure S4). This result suggests that antigen presentation by pHLIP is not prone to rapid internalization as it is the case with ligand-bound membrane receptors such as the prostate-specific membrane antigen (PSMA) (*46*). Finally, fluorescence microscopy experiments further confirmed the flow cytometry results by showing the specificity and pH-dependence of anti-DNP antibody recruitment (Figure 2B). Interestingly, the pHLIP(D25E) conjugate shows larger recruitment that the one based on pHLIP(D14Gla,D25Aad), despite the fact that its pH_50_ is expected to be lower (6.5 vs. 6.8). This difference in recruitment may be attributed to the higher helical content observed for pHLIP(D25E) than for pHLIP(D14Gla,D25Aad) (Figure S2), which may be an indication of a higher level of insertion. Measurements of tryptophan fluorescence emission, which are commonly used to monitor pHLIP insertion in lipid membranes, could not be used here because DNP is a known quencher of tryptophan. Nevertheless, because DNP-pHLIP(D25E) showed the highest level of fluorescence (i.e., the highest level of antibody recruitment) and the highest pH-selectivity among the three conjugates (Figure 2A), it was selected as the lead conjugate for further evaluation.

**Figure 2.**
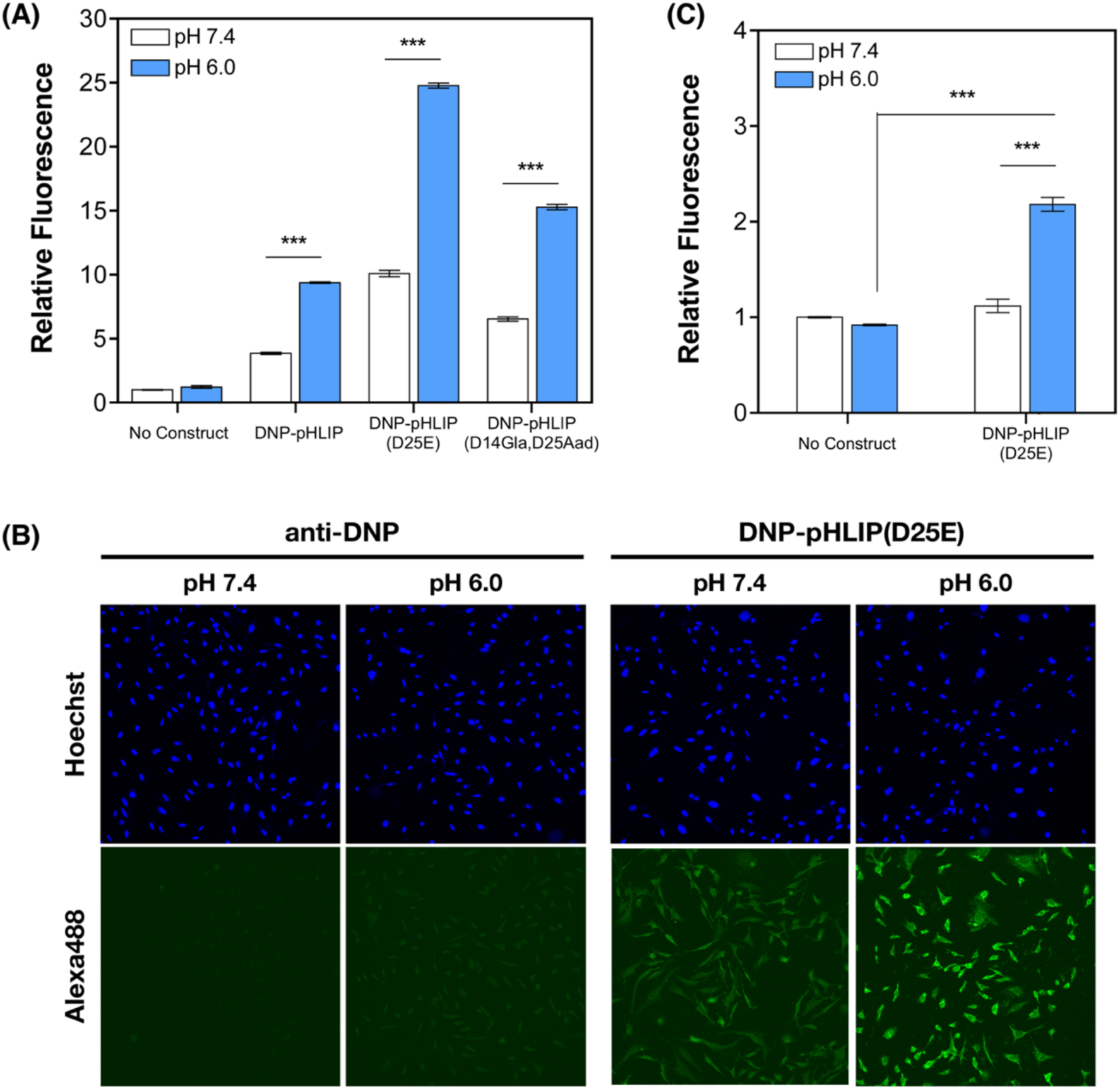
DNP-pHLIP conjugates recruits anti-DNP antibodies to the surface of cancer cells in a pH-dependent manner. **(A)** MDA-MB-231 cells were treated with 1 µM DNP-pHLIP conjugates at pH 7.4 or 6.0 and incubated with anti-DNP-A488. The amount of antibody recruitment was quantified by flow cytometry. Relative fluorescence represents the fold increase over cells incubated at pH 7.4 with anti-DNP-A488 only. **(B)** Representative immunofluorescence microscopy images of MDA-MB-231 cells treated with DNP-pHLIP(D25E), anti-DNP-A488 (green), and Hoescht (blue). **(C)** MDA-MB-231 cells were treated with 1 µM DNP-pHLIP(D25E) at pH 7.4 or 6.0 and incubated with 12.5% PHS. The amount of antibody recruitment was quantified by flow cytometry with FITC-labeled anti-human IgG. Relative fluorescence represents the fold increase over cells incubated with PHS only at pH 7.4. Results are shown as mean ± SEM (*n* = 3). Statistical significance was assessed using unpaired *t* test (at 95% confidence intervals): \*\*\**p* ≤ 0.001.

In order to better mimic physiological conditions, we determined the ability of DNP-pHLIP(D25E) to recruit anti-DNP antibodies to cancer cells directly from pooled human serum (PHS), as anti-DNP antibodies are present in normal human serum. Briefly, MDA-MB-231 cells were treated with DNP-pHLIP(D25E) at pH 7.4 or 6.0 and then incubated with 12.5% PHS instead of purified anti-DNP-A488 antibodies (*47*). Detection of anti-DNP recruitment was performed using FITC-labeled anti-human IgG antibodies. Notably, treatment of cancer cells with DNP-pHLIP(D25E) led to a significant increase in relative fluorescence at pH 6.0 compared to pH 7.4 (Figure 2C), an indication of the anti-DNP recruitment directly from serum.

### DNP-pHLIP(D25E) induces complement-dependent and antibody-dependent cellular cytotoxicity

Having established that DNP-pHLIP(D25E) promoted the recruitment of anti-DNP antibodies, we next sought to establish its ability to induce CDC and ADCC of MDA-MB-231 cells. Indeed, these processes are known to be mediated by interactions between the Fc-regions of antibodies and either the proteins of the complement cascade present in the serum or Fcγ receptors (FcγR) present on cytotoxic effector cells contained in peripheral blood (e.g., natural killer cells and macrophages). Therefore, to first determine whether DNP-pHLIP(D25E) could induce CDC of cancer cells, MDA-MB-231 cells were treated with the conjugate and incubated with anti-DNP IgG antibody prior to the addition of PHS. Cytotoxicity was assessed by lactate dehydrogenase (LDH) release after a 4-hour incubation. Cells treated with 1 µM DNP-pHLIP(D25E) at pH 6.0 showed 10% cytotoxicity, whereas treatment at pH 7.4 induced no toxicity at all (Figure 3A). While moderate, this increase in cytotoxicity is comparable with what has been observed with receptor-targeted DNP conjugates (*43*). Next, we examined whether DNP-pHLIP(D25E) could elicit killing through the ADCC pathway using isolated human peripheral blood mononuclear cells (PBMCs). Briefly, MDA-MB-231 cells were treated with varying concentrations of the conjugate at pH 7.4 and 6.0, incubated with anti-DNP antibodies, and mixed with PBMCs at an effector to target ratio of 50:1. Cell viability was determined by LDH release. Strikingly, a concentration- and pH-dependent cytotoxicity was at sub-micromolar doses of DNP-pHLIP(D25E) (Figure 3B). However, treatment with 1 µM of DNP-pHLIP(D25E) resulted in better pH-selectivity and significantly higher levels of cytotoxicity (40%) than with 0.1 µM (20%) or with PHS treatments (Figure 3A; 10%). Importantly, cytotoxicity was not observed when cells were incubated with mock anti-FITC antibodies or in the absence of antibodies prior to addition of PBMCs (Figure S5A) or when cells were treated with up to 10 µM DNP-pHLIP(D25E) alone (Figure S6). These results establish that the observed cytotoxic is mediated specifically by DNP-pHLIP(D25E) and anti-DNP antibodies, and that DNP-pHLIP(D25E) is not toxic on its own.

**Figure 3.**
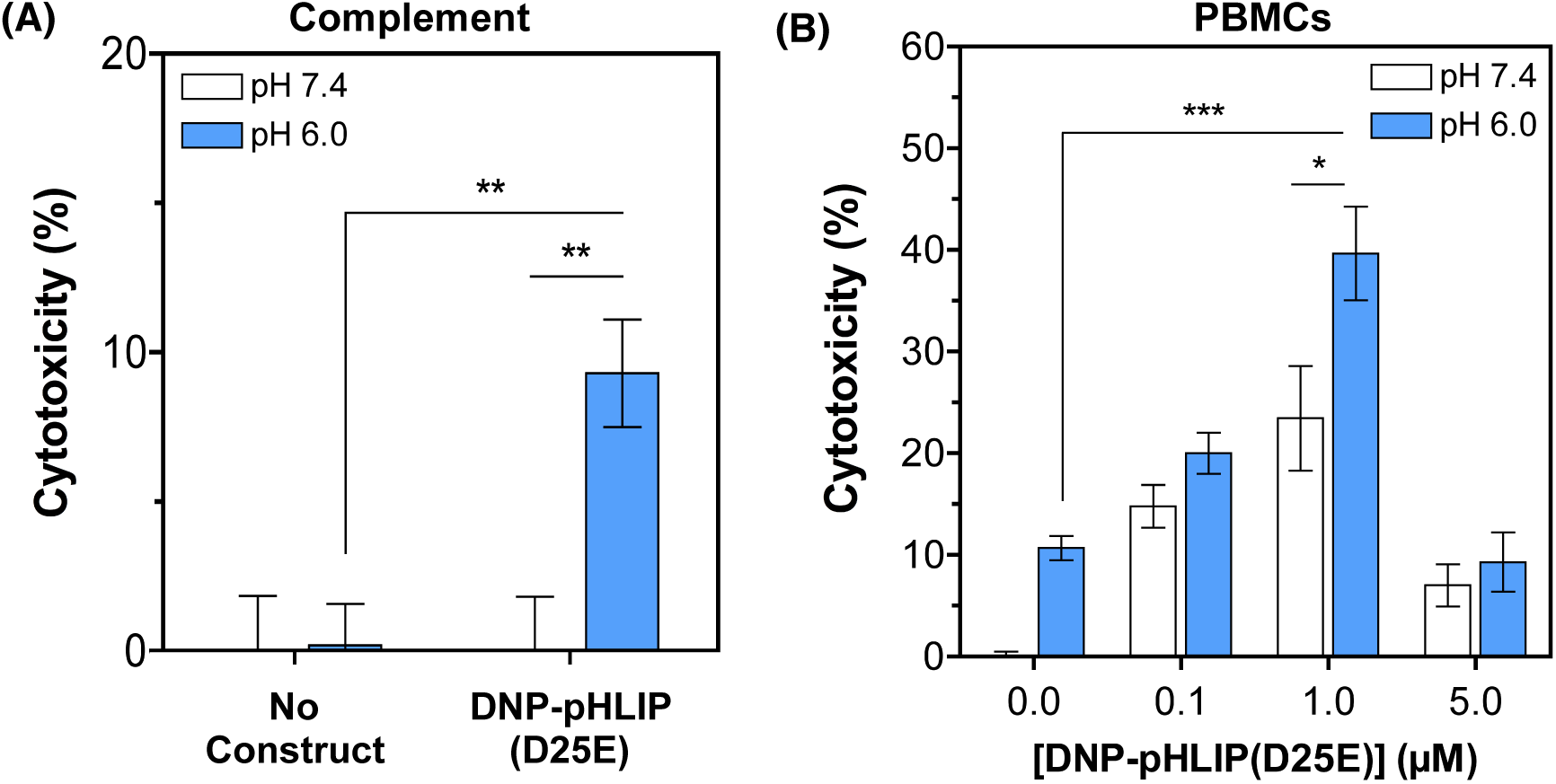
DNP-pHLIP(D25E) induces CDC and ADCC. MDA-MB-231 cells were treated with DNP-pHLIP(D25E) at pH 7.4 or 6.0 and incubated with anti-DNP antibody. LDH release was measured after 4 hours of incubation with **(A)** 12.5% pooled human serum, **(B)** PBMCs at an effector to target ratio of 50:1 Cytotoxicity was calculated by percent difference in LDH release between cells treated with anti-DNP antibody only at pH 7.4 and cells treated with DNP-pHLIP(D25E). Results are shown as mean ± SEM (*n* = 3-12). Statistical significance was assessed using unpaired *t* test (at 95% confidence intervals): \*\*\**p* ≤ 0.001, \*\**p* ≤ 0.01, \**p* ≤ 0.05.

Finally, in addition to induce killing via PBMCs, we sought to evaluate whether DNP-pHLIP(D25E) could activate engineered natural killer (NK) cells. There has been recently tremendous enthusiasm for engineered NK cells as a source of “off-the-shelf” allogeneic NK supplementation. More specifically, an engineered immortal NK-92 cell line that expresses a variant of the CD16 receptor (FcγRIII) with high affinity to the Fc-domain of IgG1 isotype (NK-92-CD16^+^ or haNK cells) could be widely applicable to cancer immunotherapies (*48-51*). These particular cells are readily expanded and cryopreserved for administration into any patient (no toxicity has been observed in human clinical trials) to increase the density of NK cells that engage tightly with Fc-domains. NK-92-CD16^+^ cells have many promising features including: lack of graft-versus-host disease, superior cytotoxicity towards target cells, established safety in humans at high infusion doses, greatly reduced cost due to facile culture-based cell expansion of a single universal cell line, and once irradiated prior to injection they do not expand *in vivo* (in contrast to CAR-T cells). Thanks to their low toxicity and potential universality, haNK cells are under clinical evaluation (*52-57*) to potentiate various cancer immunotherapeutic agents. Similar to experiments using PBMCs, MDA-MB-231 cells were first treated with increasing concentrations of DNP-pHLIP(D25E), and then mixed with NK-92-CD16^+^ cells at an effector to target ratio of 5:1 (Figure 4). pH-dependent cytotoxic effect was observed when the cells were treated with concentrations ranging from 0.1 to 5 µM, with maximum cytotoxicity and pH-selectivity at 500 nM (∼40%; Figure 4). Similar to what was observed with PBMCs, cytotoxicity was not observed when cells were incubated with mock anti-FITC antibodies or in the absence of antibodies before the addition of NK-92-CD16^+^ cells (Figure S5B), a clear indication that the observed cytotoxic is specifically mediated by DNP-pHLIP(D25E). Interestingly, a bell-shaped response to the concentration of DNP-pHLIP(D25E) was observed in cell killing experiments with both PBMCs (Figure 3B) and NK-92-CD16^+^ (Figure 4). This distinct pattern, also known as the “hook effect”, is frequently observed experimentally with bifunctional agents as a result of unique binding dynamics (*12, 58, 59*). At high concentrations, unbound compound can prevent cytotoxic effects due to univalent saturation thereby preventing bivalent bridging of effector and target cells (*60*).

**Figure 4.**
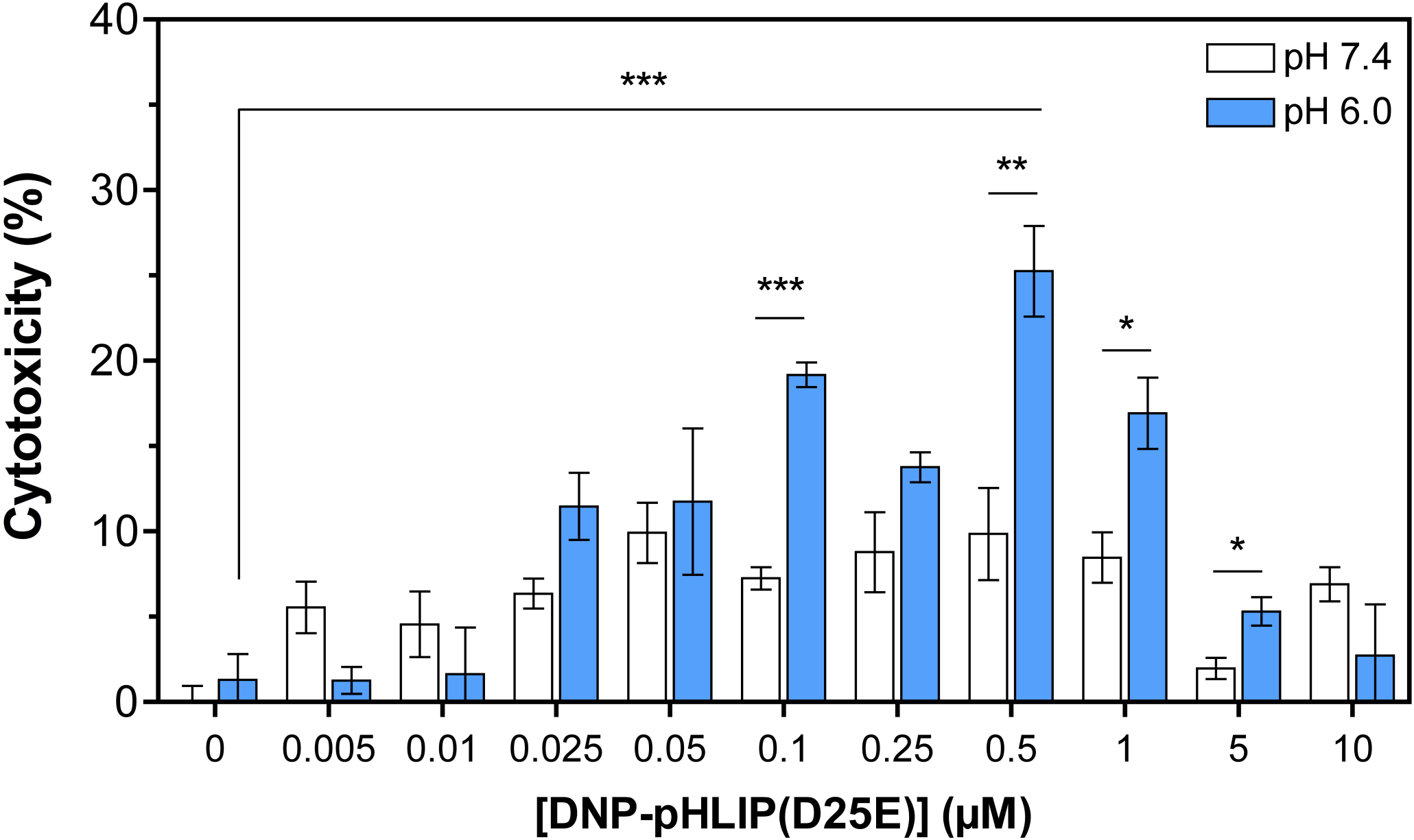
DNP-pHLIP(D25E) induces ADCC through engineered NK cells. MDA-MB-231 cells were treated with DNP-pHLIP(D25E) at pH 7.4 or 6.0 and incubated with anti-DNP antibody. LDH release was measured after 4 hours of incubation with NK-92-CD16^+^ cells at an effector to target ratio of 5:1. Cytotoxicity was calculated by percent difference in LDH release between cells treated with anti-DNP antibody only at pH 7.4 and cells treated with DNP-pHLIP(D25E). Results are shown as mean ± SEM (*n* = 3-12). Statistical significance was assessed using unpaired *t* test (at 95% confidence intervals): \*\*\**p* ≤ 0.001, \*\**p* ≤ 0.01, \**p* ≤ 0.05.

## Conclusion

In conclusion, we have designed and tested a novel class of immunotherapeutic agents that exploits the acidic microenvironment of tumors to selectively decorate the surface of cancer cells with immunogenic epitopes. We showed that the most potent agent triggers opsonization and induces CDC- and ADCC-based killing of cancer cells. Most significantly, this lead agent induced selective cell death *via* an engineered NK cell line, opening the possibility of allogeneic NK supplementation without the need of pre-vaccination. Optimization efforts and evaluation of this strategy in animal models, as well as the investigation of how it can be used to graft exogenous antigens are currently ongoing in our laboratories.

## Methods

### Solid-phase peptide synthesis

All pHLIP variants were prepared by Fmoc solid-phase chemistry with a CEM Liberty Blue microwave peptide synthesizer using rink amide resin (CEM, 0.19 mmol/g loading capacity). To prepare DNP-pHLIP conjugates, N-Fmoc-N-2,4-dinitrophenyl-L-lysine (Chem-Impex #05734) was coupled to the N-terminus of each pHLIP variant on resin. Peptides were purified via reverse-phase high-performance liquid chromatography (RP-HPLC; Phenomenex Luna Omega 5 µm 250 × 21.2 mm C18; flow rate 5 mL/min; phase A: water 0.1% TFA; phase B: acetonitrile 0.1% TFA; gradient 60 min from 95/5 A/B to 0/100 A/B). The purity of the peptides was determined by RP-HPLC (Phenomenex Luna Omega 5 µm 250 × 10 mm C18; flow rate 5 mL/min; phase A: water 0.01% TFA; phase B: acetonitrile 0.01% TFA; gradient 60 min from 95/5 A/B to 0/100 A/B), and their identity was confirmed via matrix-assisted laser desorption/ionization time of flight (MALDI-TOF) mass spectroscopy (Shimadzu 8020).

Sequence of peptides used in this study:

DNP-pHLIP(WT):

(DNP)-KEQNPIYWARYA<u>D</u>WLFTTPLLLL<u>D</u>LALLVDADEGTG

DNP-pHLIP(D25E):

(DNP)KEQNPIYWARYADWLFTTPLLLL<u>E</u>LALLVDADEGTG

DNP-pHLIP(D14GlaD25Aad):

(DNP)-KEQNPIYWARYA<u>Gla</u>WLFTTPLLLL<u>Aad</u>LALLVDADEGTG

### Sample preparation of CD measurements

All peptide constructs were solubilized to 20 µM in 5 mM sodium phosphate, pH 8.0. Each construct was diluted to a final concentration of 7 μM before analysis. For measurements in vesicles, 1-palmitoyl-2-oleoyl-sn-glycero-3-phosphocholine (POPC) was dried as a thin film and held under vacuum for at least 24 hours. The lipids were rehydrated in 5 mM sodium phosphate, pH 8.0, for at least 30 min with periodic gentle vortexing. The resulting large multilamellar vesicles were freeze-thawed for seven cycles and subsequently extruded through a polycarbonate membrane with 100 nm pores using a Mini-Extruder (Avanti Polar Lipids) to produce large unilamellar vesicles (LUVs). The constructs were incubated with the resulting LUVs at a 1:300 ratio. The pH was adjusted to the desired experimental values with HCl, and the samples were incubated at room temperature for 30 minutes prior to spectroscopic analysis.

### CD spectroscopy

Far-UV CD spectra were recorded on a Jasco J-815 CD spectrometer equipped with a Peltier thermal-controlled cuvette holder (Jasco). Measurements were performed in 0.1 cm quartz cuvette. CD intensities are expressed in mean residue molar ellipticity [θ] calculated from the following equation:

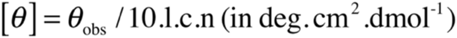

where, θ_obs_ is the observed ellipticity in millidegrees, l is the optical path length in centimeters, c is the final molar concentration of the peptides, and n is the number of amino acid residues. Raw data was acquired from 260 to 200 nm at 1 nm intervals with a 100 nm/min scan rate, and at least five scans were averaged for each sample. The spectrum of POPC liposomes was subtracted out from all construct samples.

### Cell culture

Human breast adenocarcinoma MDA-MB-231 were cultured in Dulbecco’s Modified Eagle’s medium (DMEM) high glucose supplemented with 10% fetal bovine serum (FBS), 100 units/ml penicillin and 0.1 mg/ml streptomycin. CD16^+^ natural killer NK-92 cells (ATCC PTA-6967) were cultured in minimum essential medium (MEMα) without nucleosides supplemented with 0.1 mM 2-mercaptoethanol, 0.2 mM inositol, 0.02 mM folic acid, 12.5% FBS, 12.5% horse serum, 100-200 units/mL interleukin-2 (IL-2), 100 units/mL penicillin, and 0.1 mg/mL streptomycin. Cells were cultured in a humidified atmosphere of 5% CO_2_ at 37 °C.

### Antibody recruitment assay

Peptide constructs were resuspended in 5 mM sodium phosphate, pH 9.0, to a concentration of 20 µM and incubated at room temperature (RT) for an hour. Immediately following the incubation, the peptide was diluted to 12.5 µM by addition of PBS pH 7.4. The peptide was further diluted to the appropriate concentration so that upon pH adjustment the desired treatment concentration (1 µM) is obtained with 1.6:1 5 mM sodium phosphate pH 9.0:PBS pH 7.4. MDA-MB-231 cells were harvested and washed twice with PBS pH 7.4. Next, 300,000 cells were treated in suspension with 1 µM peptide constructs for 5 minutes at 37 °C at pH 7.4 or 6.0. After treatment, the cells were washed once with PBS at the same pH as treatment with 10% FBS. For experiments with DNP-pHLIP conjugates, the cells were subsequently incubated with 25 µg/mL of Alexa Fluor 488 conjugated anti-dinitrophenyl polyclonal rabbit antibody (anti-DNP-A488, Invitrogen #A11097) in PBS pH 7.4 with 10% FBS for 30 minutes at 4 °C. For control experiments with DNP-pHLIP conjugates, the cells were incubated in PBS pH 7.4 with 10% FBS with and without 25 µg/mL FITC-labeled anti-human IgG antibody (Sigma F9512) for 30 minutes at 4 °C. The cells were washed once with PBS at the same pH as treatment with 10% FBS and fixed with 4% paraformaldehyde (PFA) for 10 minutes at 4 °C. The cells were resuspended in PBS and analyzed by flow cytometry using a BDFacs Canto II flow cytometer equipped with a 488 nm argon laser and a 530/30 bandpass filter. A minimum of 3,000 events were counted for each experimental condition. The data was analyzed using FACSDiva version 6.1.1 software. The fluorescence data are expressed as mean arbitrary fluorescence units and were gated to include all healthy mammalian cells.

### Immunofluorescence microscopy

MDA-MB-231 cells were seeded on 22 mm x 22 mm chambered coverslips pretreated with poly-l-lysine to be ∼70% confluent after 16 hours. DNP-pHLIP(D25E) was solubilized as described in the antibody recruitment assay. The cells were treated with 1 µM DNP-pHLIP(D25E) for 5 minutes at 37°C at pH 7.4 or pH 6.0. The cells were washed once with PBS at the same treatment pH, fixed with ice cold methanol for 10 minutes, and washed two more times. The coverslips were incubated for 1 hour at 37 °C with Alexa Fluor 488 conjugated anti-dinitrophenyl-KLH polyclonal rabbit antibody (anti-DNP-A488, Invitrogen #A11097) at 50 µg/mL in PBS. After five washes with PBS, the cells were stained with Hoechst (Invitrogen #H3570). The coverslips were mounted on slides with Fluoromount (Sigma-Aldrich #F4680) before being imaged by a Nikon Eclipse Ti microscope with a 20x objective.

### Antibody recruitment from pooled human serum

DNP-pHLIP(D25E) was prepared as described for the antibody recruitment assay. MDA-MB-231 cells were harvested and washed twice with PBS pH 7.4. Next, 300,000 cells were treated in suspension with 1 µM DNP-pHLIP(D25E) for 5 minutes at 37 °C at pH 7.4 or 6.0. After treatment, the cells were washed once with PBS at the same pH as treatment with 10% FBS. The cells were subsequently incubated with human IgG isotype control (Invitrogen #02-7102) at 100 µg/mL in PBS with 1% BSA for 20 minutes at 4 °C. The cells were washed once with PBS at the same pH as treatment with 10% FBS and then incubated with 12.5% pooled human complement serum (Innovative Research, Inc. #IPLA-CSER-22267) in PBS pH 7.4 at 4 °C for 20 minutes. The cells were washed once with PBS at the same pH as treatment with 10% FBS and incubated with 25 µg/mL FITC-labeled anti-human IgG antibody (Sigma F9512) in PBS pH 7.4 with 10% FBS for 30 minutes at 4 °C. After washing, the cells were fixed with 4% PFA for 10 minutes at 4 °C, and immediately analyzed by flow cytometry as previously described.

### Complement-dependent cytotoxicity

DNP-pHLIP(D25E) was prepared as described for the antibody recruitment assay. MDA-MB-231 cells were harvested and washed twice with PBS pH 7.4. Next, 240,000 cells were treated in suspension with 1 µM DNP-pHLIP(D25E) for 5 minutes at 37 °C at pH 7.4 or 6.0. After treatment, the cells were washed once with PBS at the same pH as treatment with 10% FBS and then incubated with 25 µg/mL unconjugated anti-DNP-KLH polyclonal rabbit antibody (Invitrogen #A6403) in PBS pH 7.4 with 10% FBS for 30 minutes at 4 °C. Immediately following antibody incubation, pooled human complement serum (Innovative Research, Inc. #IPLA-CSER-22267) was added to 12.5% in DMEM and incubated for an additional 4 hours at 37 °C. The cells were harvested, the supernatant was collected, and lysis was quantified by lactate dehydrogenase (LDH) release. Following the manufacturer’s protocol (ThermoFisher #88953), the cell media supernatant was transferred to a 96 well plate in triplicate and an equal volume of LDH reaction mixture was added to each well. The plate was incubated for 30 minutes at room temperature. The absorbance was read at 490 nm and 680 nm on an Infinite® 200 PRO Plate Reader (Tecan). Cytotoxicity was calculated by: cytotoxicity (%) = (LDH release – normal release)/(normal release) x 100%. Normal release is defined by cells treated with anti-DNP-KLH polyclonal rabbit antibody but no peptide construct.

### Isolation of peripheral blood mononuclear cells

Human blood from a single donor was mixed with EasySep buffer (StemCell Technologies), layered over SepMate-50 conical tubes (StemCell Technologies) prepared with Lymphoprep™ density gradient medium (StemCell Technologies), and centrifuged according to manufacturer’s instructions. Isolated PBMCs were mixed with Muse Count and Viability Reagent (EMD Millipore), and then counted using a Muse Cell Analyzer. PBMCs were cryopreserved using the CTL-Cryo™ ABC Media Kit (ImmunoSpot), following manufacturer’s protocol. The resulting suspension was aliquoted into cryovials and placed in a -80 °C freezer. After 4-16 hours, the cryovials were transferred to liquid nitrogen storage.

### Antibody-dependent cellular cytotoxicity

DNP-pHLIP(D25E) was prepared as previously described for the antibody recruitment assay. MDA-MB-231 cells were harvested and washed twice with PBS pH 7.4. Next, 150,000 cells were treated in suspension with DNP-pHLIP(D25E) for 5 minutes at 37 °C at pH 7.4 or 6.0. The cells were washed once at 4 °C with PBS at the same pH as treatment with 10% FBS and then incubated with 0.025 mg/mL unconjugated anti-DNP-KLH polyclonal rabbit antibody (Invitrogen #A6403) in PBS with 10% FBS for 30 minutes at 4 °C. Following antibody incubation, the solution was removed and the cells were resuspended in human Peripheral Blood Mononuclear Cells (PBMCs) at a 50:1 effector to target ratio in Roswell Park Memorial Institute (RPMI) 1640 medium or NK-92 cells CD16^+^ natural killer NK-92 cells (ATCC, PTA-6967) in MEMα at a 5:1 effector to target ratio in MEMα, and incubated for 4 hours at 37 °C. The cells were harvested, the supernatant was collected, and lysis was quantified by LDH release as previously described for CDC. Cytotoxicity was calculated by: cytotoxicity (%) = (LDH release – normal release)/(normal release) x 100%. Normal release is defined by target cells treated with anti-DNP-KLH polyclonal rabbit antibody and effector cells but no peptide construct. For control experiments, normal release is defined by target cells treated with effector cells only or target cells only treated at pH 7.4.

## Acknowledgements

This work was supported by the National Cancer Institute [grant number R21CA181868] to D.T. and by the National Institute of General Medical Sciences [grant number R35GM124893-01] to M.M.P.; and internal funds from Lehigh University to D.T. and M.M.P.

## Author Contributions

D.T. and M.M.P designed research; J.W., E.L.S., E.B., M.S.F, N.J.F and T.B. performed research; J.W., E.L.S., E.B., M.S.F, M.M.P and D.T. analyzed the data; J.W., E.L.S., M.M.P and D.T. wrote the paper.

## Abbreviations

ADCC: antibody-dependent cellular cytotoxicity.
CD: Circular Dichroism.
CDC: complement-dependent cytotoxicity.
DNP: 2,4-dinitrophenol.
LDH: lactate dehydrogenase.
MALDI-TOF: Matrix-assisted laser desorption/ionization time of flight.
PBMC: peripheral blood mononuclear cells.
pHLIP: pH(Low) Insertion Peptide.
POPC: 1-palmitoyl-2-oleoyl-sn-glycero-3-phosphocholine.
RP-HPLC: reverse-phase high-performance liquid chromatography.
WT: Wild-type.

