## Supplementary Material for "pH-Dependent Grafting of Cancer Cells with Antigenic Epitopes Promotes Selective Antibody-Mediated Cytotoxicity"

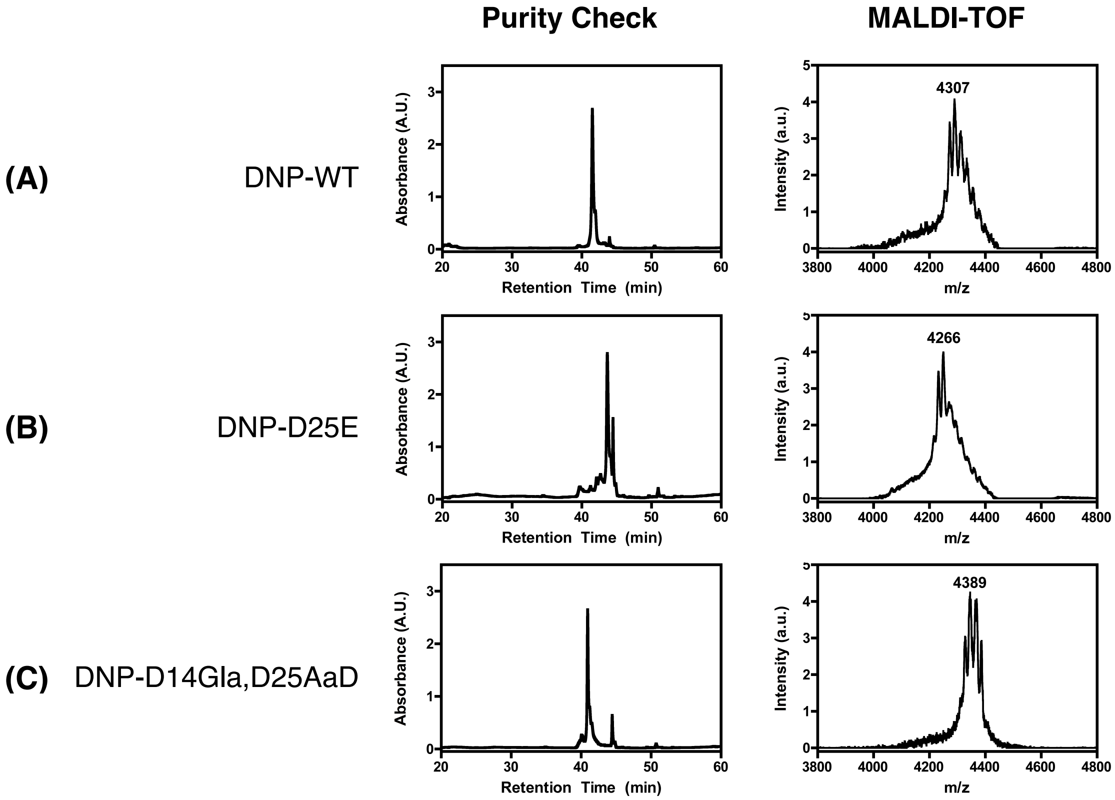


**Figure S1. Purity check by RP-HPLC and MALDI-TOF MS spectra of DNP-pHLIP conjugates. (A)** DNP-pHLIP: purity >99%; calculated ($MH^+$) = 4306, found ($MH^+$) = 4307 **(B)** DNP-pHLIP (D25E): purity >90%; calculated ($MH^+$) = 4263, found ($MH^+$) = 4266 **(C)** DNP-pHLIP(D14Gla,D25AaD): purity >95%; calculated ($MH^+$) = 4389, found ($MH^+$) = 4389.

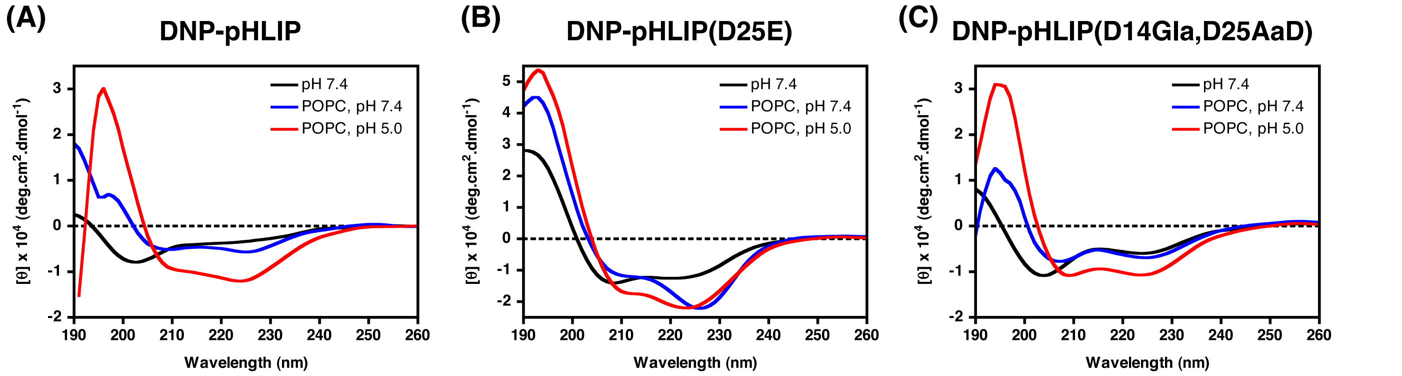


**Figure S2. Circular dichroism spectra of DNP-pHLIP conjugates. (A)** DNP-pHLIP, **(B)** DNP-pHLIP(D25E), and **(C)** DNP-pHLIP(D14Gla,D25Aad) in the absence (black) or presence of large unilamellar POPC lipid vesicles at pH 7.4 (blue) and pH 5.0 (red).

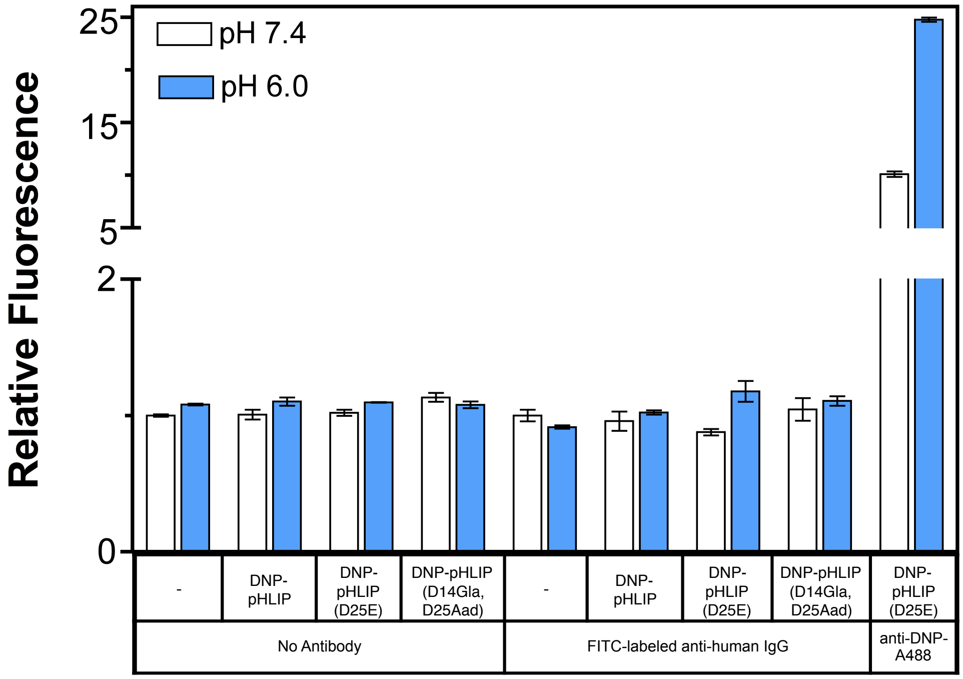


**Figure S3. Observed increase in fluorescence results from specific recruitment of antibodies by DNP-pHLIP conjugates.** MDA-MB-231 cells were treated with DNP-pHLIP conjugates at 1 µM at pH 7.4 or 6.0 and incubated with no antibody or FITC-labeled anti-human IgG. Antibody recruitment was analyzed by flow cytometry. Relative fluorescence represents the fold increase over cells incubated at pH 7.4 only (no peptide or antibody treatment). Results are shown as mean ± SEM ($n = 3$).

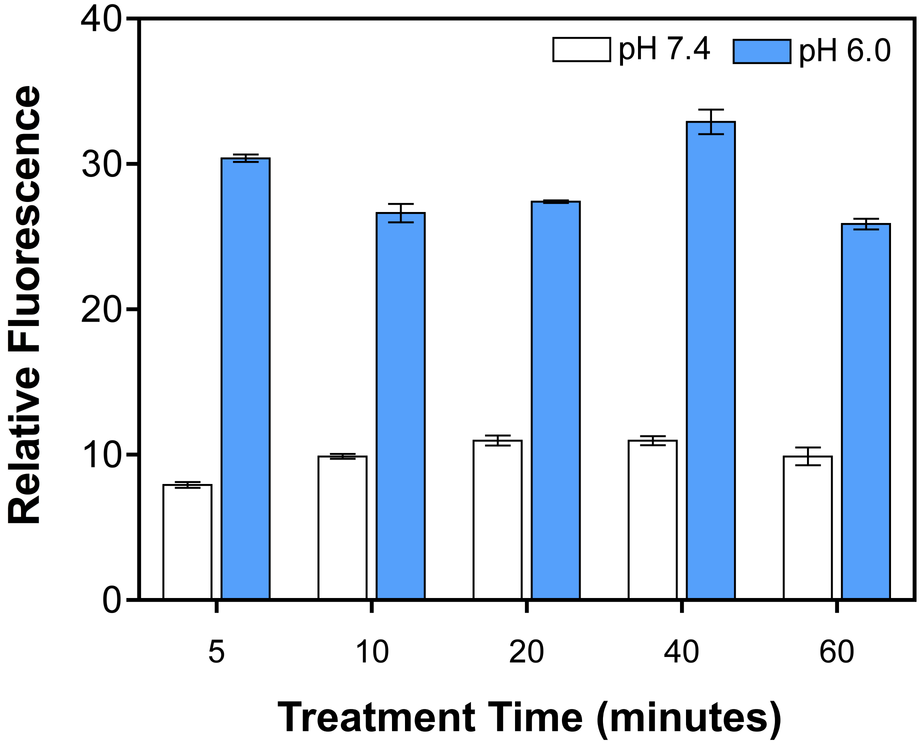


**Figure S4. DNP-pHLIP(D25E) remains at the surface of cells over an hour incubation.** MDA-MB-231 cells were treated with 1 μM DNP-pHLIP(D25E) at pH 7.4 or 6.0 for the indicated times at 37 ˚C and incubated with anti-DNP-A488 for 30 minutes. Antibody recruitment was analyzed by flow cytometry. Relative fluorescence represents the fold increase over cells incubated with anti-DNP-A488 only at pH 7.4. Results are shown as mean ± SEM ($n$ = 3).

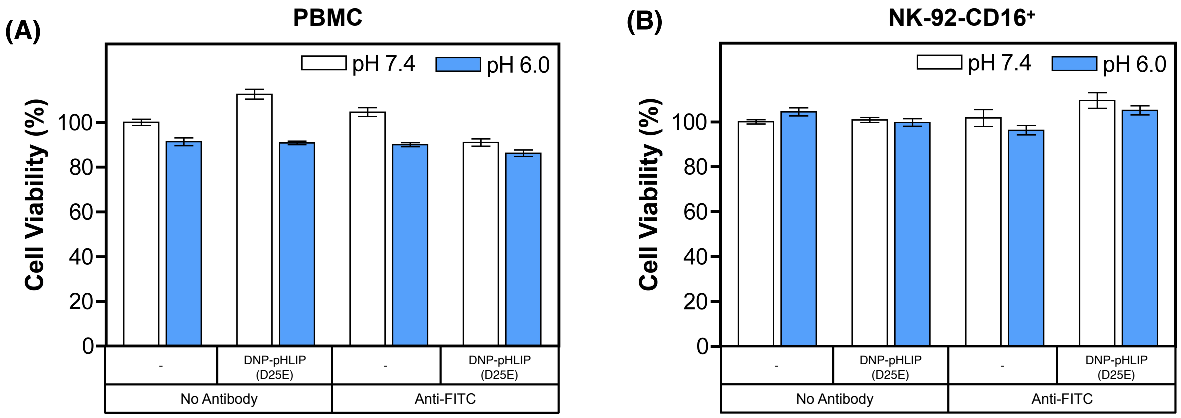


**Figure S5. DNP-pHLIP(D25E)-mediated cytotoxicity is antibody-dependent.** MDA-MB-231 cells were treated with 1 or 0.5 µM DNP-pHLIP(D25E) at pH 7.4 or 6.0 and incubated with anti-FITC polyclonal rabbit antibody for 30 minutes. LDH release was measured after 4 hours of incubation with **(A)** PBMCs at an effector to target ratio of 50:1 and **(B)** NK-92-CD16+ cells at an effector to target ratio of 5:1. All measurements were normalized to cells incubated at pH 7.4 only (no peptide or antibody treatment) as 100% cell viability. Results are shown as mean ± SEM ($n$ = 3).

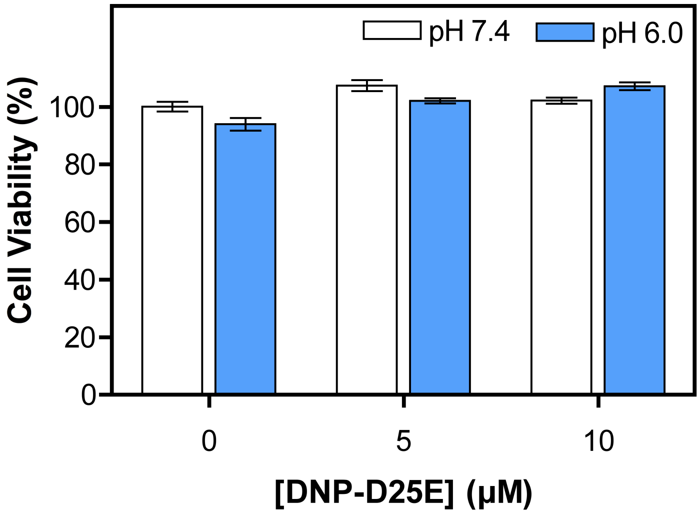


**Figure S6. DNP-pHLIP(D25E) is not toxic to cells.** MDA-MB-231 cells were treated with DNP-pHLIP(D25E) at pH 7.4 or 6.0, and cell viability was assessed by LDH release after 4 hours of incubation. All measurements were normalized to cells incubated at pH 7.4 only (no peptide or antibody treatment) as 100% cell viability. Results are shown as mean ± SEM ($n$ = 3).
